# Stronger effect of temperature on body growth in cool than in warm populations suggests lack of local adaptation

**DOI:** 10.1101/2024.01.17.575983

**Authors:** Max Lindmark, Jan Ohlberger, Anna Gårdmark

## Abstract

Body size is a key functional trait that has declined in many biological communities, partly due to changes in individual growth rates in response to climate warming. However, our understanding of growth responses in natural ecosystems is limited by relatively short time series without large temperature contrasts and unknown levels of adaptation to local temperatures across populations. In this study, we collated back-calculated length-at-age data for the fish Eurasian perch (*Perca fluviatilis*) from 10 populations along the Baltic Sea coast between 1953–2015 (142023 length-at-age measurements). We fitted individual-level growth trajectories using the von Bertalanffy growth equation, and reconstructed local temperature time series using generalized additive models fitted to three data sources. Leveraging a uniquely large temperature contrast due to climate change and artificial heating, we then estimated population-specific and global growth-temperature relationships using Bayesian mixed models, and evaluated if they conformed to local adaption or not. We found little evidence for local adaptation in the temperature-dependence of individual growth curves. Instead, population-specific curves mapped onto a global curve, resulting in body growth increasing with warming in cold populations but decreasing in warm populations. Understanding to which degree the effects of warming on growth and size are population-specific is critical for generalizing predictions of climate impacts on growth, which is a key biological trait affecting multiple levels of biological organisation from individuals to ecosystem functioning.

## Introduction

Body growth is an individual-level trait that is relevant to ecology across all levels of biological organisation (Peters 1983, Barneche *et al*. 2019). In aquatic systems in particular, body growth is sensitive to environmental conditions, is related to individual fitness (Sibly *et al*. 2018), determines species interactions and dictates how much energy is transferred between trophic levels (Lindeman 1942). It is also directly related to body size, which is a key ecological trait (Peters 1983) that is correlated with diet, survival and reproductive success (Barneche *et al*. 2018) and largely shapes size-dependent species interactions (Ursin 1973, Werner and Gilliam 1984).

In ectotherms such as fish, environmental temperature has a large influence on body growth via the effects on metabolic rate (Jobling 1997, Brown *et al*. 2004). For species living at temperatures cooler than that which maximizes growth, as commonly observed (Lindmark *et al*. 2022, Tewksbury *et al*. 2008), a slight increase in temperature is likely to be beneficial to growth. Body growth or size-at-age of fish in natural environments, is commonly observed to correlate positively with temperature, especially for small or young fish (Thresher *et al*. 2007, Baudron *et al*. 2014, Huss *et al*. 2019, Oke *et al*. 2022, Lindmark *et al*. 2023). The effects on old fish, however, often are smaller or negative (Morrongiello *et al*. 2014, van Dorst *et al*. 2019, Ikpewe *et al*. 2020), although there are exceptions (Lindmark *et al*. 2023) and responses can vary within populations, e.g., with sex (van Dorst *et al*. 2023). Experimental and modelling studies have pointed to that size-dependent responses of growth and size could be due to optimum growth temperatures being lower for larger fish (Lindmark *et al*. 2022), or that warming is linked to earlier maturation, after which energy is increasingly allocated to reproduction over somatic growth (Wootton *et al*. 2022, Niu *et al*. 2023), or both (Audzijonyte *et al*. 2022). In natural systems, other factors such as competition and food limitation also influence growth directly (Oke *et al*. 2020, Cline *et al*. 2019, Ohlberger *et al*. 2023), and indirectly by reducing the optimal growth temperatures (Brett *et al*. 1969, Brett 1971, Huey and Kingsolver 2019). To understand fish responses to changing temperatures, it is therefore important to evaluate growth-temperature relationships in natural systems, and across gradients of environmental temperature.

The ability to quantify the impacts of temperature change on growth and size, or other ecological traits, is often limited by relatively short time series that contain small temperature contrasts (White 2019, Freshwater *et al*. 2023). As an alternative, studies often use space-for-time approaches (Morrongiello *et al*. 2014, van Dorst *et al*. 2019, van Denderen *et al*. 2020) to estimate the effects of temperature on growth. However, it is difficult to know to what extent we can infer effects of warming in a given location from the temperature effects estimated across locations over a limited time (Perret *et al*. 2024). Both the estimates (van Denderen *et al*. 2020) and the form of the growth-temperature relationship may differ. For example, responses to warming tend to be unimodal, whereas they can be more linear or exponential across all populations of a species (van Denderen *et al*. 2020). For projecting impacts of warming at the species level, another missing piece is to understand the extent of local adaptation to the experienced thermal environments (Eliason *et al*. 2011). That is, to what extent populations conform to a global, species-wide thermal performance curve, versus having developed local thermal response curves with local temperature optima in order to have higher fitness in their local habitats. In other taxa, such as kelp (Britton *et al*. 2024), corals (Howells *et al*. 2013), invertebrates (Sanford and Kelly 2011), and phytoplankton (Thomas *et al*. 2012), there is a growing evidence that local adaptation has lead to populations exhibiting different responses in growth (individual or population) to ocean warming. However, studies on fishes are scarcer (but see Neuheimer and Grønkjær (2012) and Beaudry-Sylvestre *et al*. (2024)). Testing this requires time series with large temperature contrasts both within and between multiple populations in the wild.

Here, we seek to understand how climate warming has affected the growth of fish across multiple populations, using *Perca fluviatilis*, hereafter perch, as a case study. Perch is a widely distributed freshwater fish, common along the Swedish Baltic Sea coast, that is not commercially exploited and has a stationary lifestyle. These characteristics make it an ideal species for analyzing effects of temperature change on growth across environmental gradients. Specifically, we quantify growth-temperature relationships from 10 populations and evaluate if there is support for site-specific growth-temperature relationships, and whether or not those are consistent with local adaptation (Fig. 1). To address this question, we collated size-at-age data from back-calculated growth-trajectories for 23 605 individual fish over seven decades spanning large temperature contrasts due to climate warming and artificial heating from nuclear power plants, and fit statistical models relating cohort-specific growth estimates to reconstructed temperatures.

**Figure 1:**
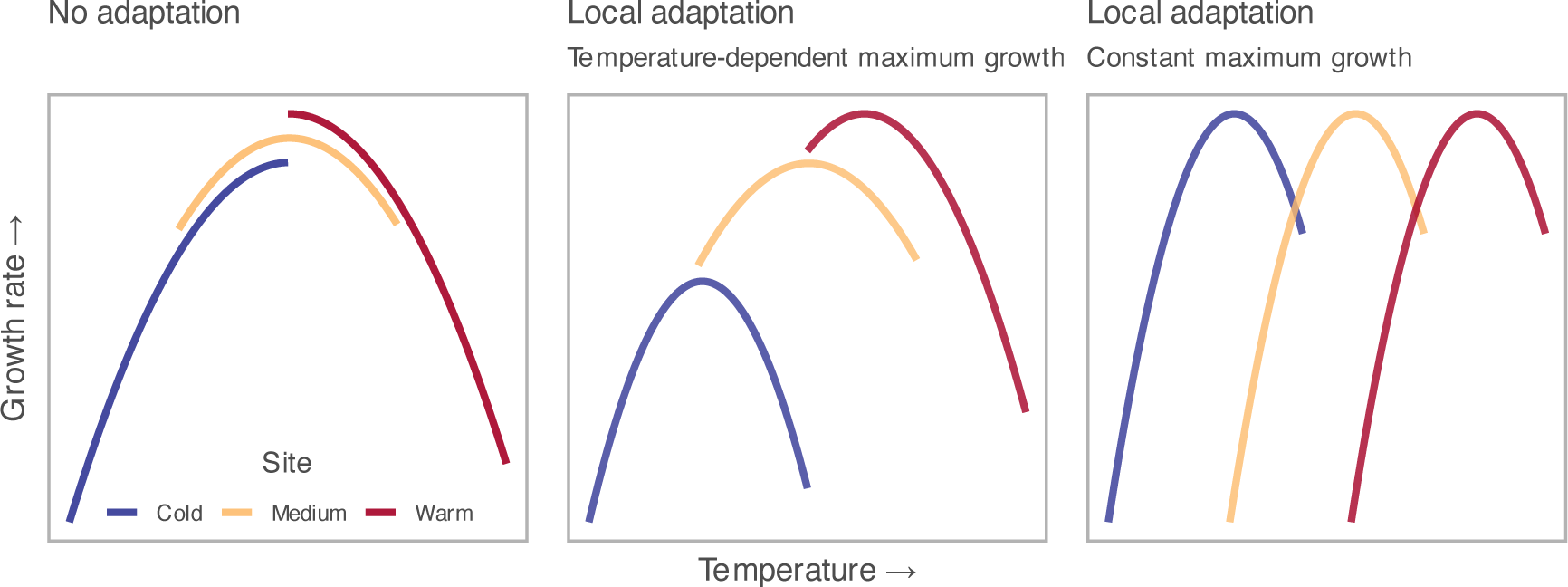
Three examples of hypothetical growth-temperature relationships across isolated populations along a temperature gradient. In all cases, the average growth rates increases with temperature across populations based on metabolic theory. In the left column, there is no local adaptation in optimum growth temperatures, such that growth in cooler (blue) populations increases with warming whereas warmer populations (yellow, red) are the first to show growth declines with warming. Hence, the effect of further warming depends on the populations distance from the global rather than the local optimum. In the middle and right columns, there is local adaptation (with maximum growth rate increasing (middle) or being constant (right) with temperature across populations), such that responses to warming are similar despite populations experiencing different temperatures.

## Methods

### Data

We compiled individual-level size-at-age data from perch and sea surface temperature data from 10 sites along the Swedish Baltic Sea coast. The longest time series started in 1953 and the shortest in 1985, and the average time series length was 34 years, which can be compared to an average generation time of ∼6 years (Froese and Pauly 2010) (Fig. 2). The temperature contrast in this data set is great both within each site and across sites (Fig. 3), due to long time series and a large latitudinal gradient. Also contributing to the large temperature range is the inclusion of sites artificially heated by warm water discharge from nearby nuclear power plants (sites (SI HA and BT in Fig. 3). The size-at-age data include information on age (at catch), total length (at catch, in millimetres), sex, and back-calculated length-at-age (in millimetres). Back-calculated length- at-age was derived from annuli rings on the operculum bones (part of the gill lid), with control counts of age done on otoliths (ear stones). This method is common in fisheries (Morrongiello and Thresher 2015, Essington *et al*. 2022), and is based on an assumed power-law relationship between the distance of annuli rings and fish length (Thoresson 1996), which allows reconstruction of the individual’s body length at each age when annuli rings where formed. Individual-level data originate from different fish monitoring programs using gill-nets. Individuals sampled for age and growth were selected from the total catch from the gill net survey in each site using random or length-stratified sub-sampling of the catch, but information on stratification method could not be retrieved for all data.

**Figure 2:**
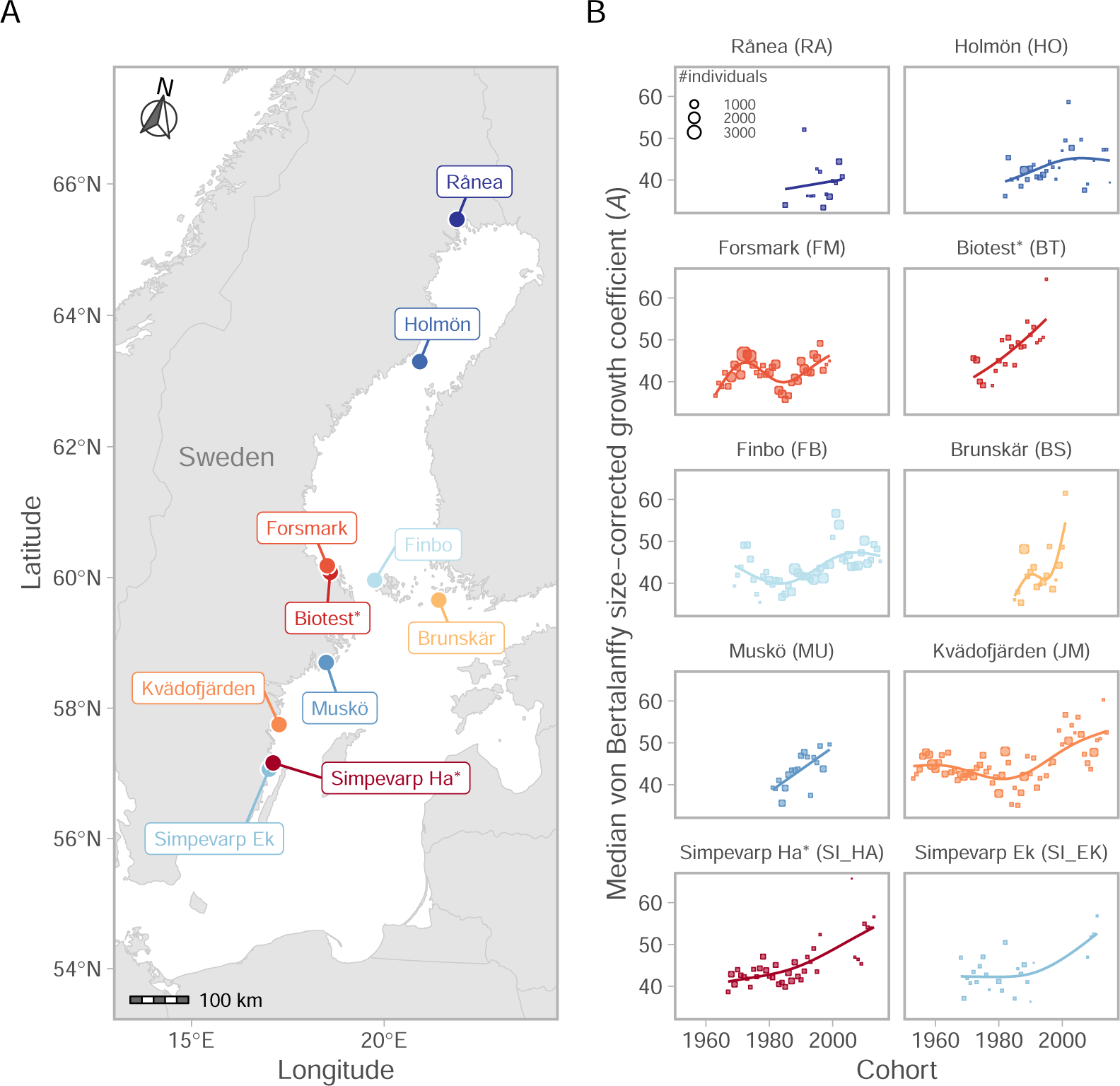
Map of sampling locations (left) and time series of the median von Bertalanffy growth coefficients by cohort (right), where colours are assigned based on the minimum temperature in the growth time series, ranging from blue (coldest) to red (warmest). Circle size corresponds to the number of individuals in that cohort and site. The lines depict fits from Generalized additive models(GAMs) with Gaussian error and basis dimension k = 5, to highlight trends.

**Figure 3:**
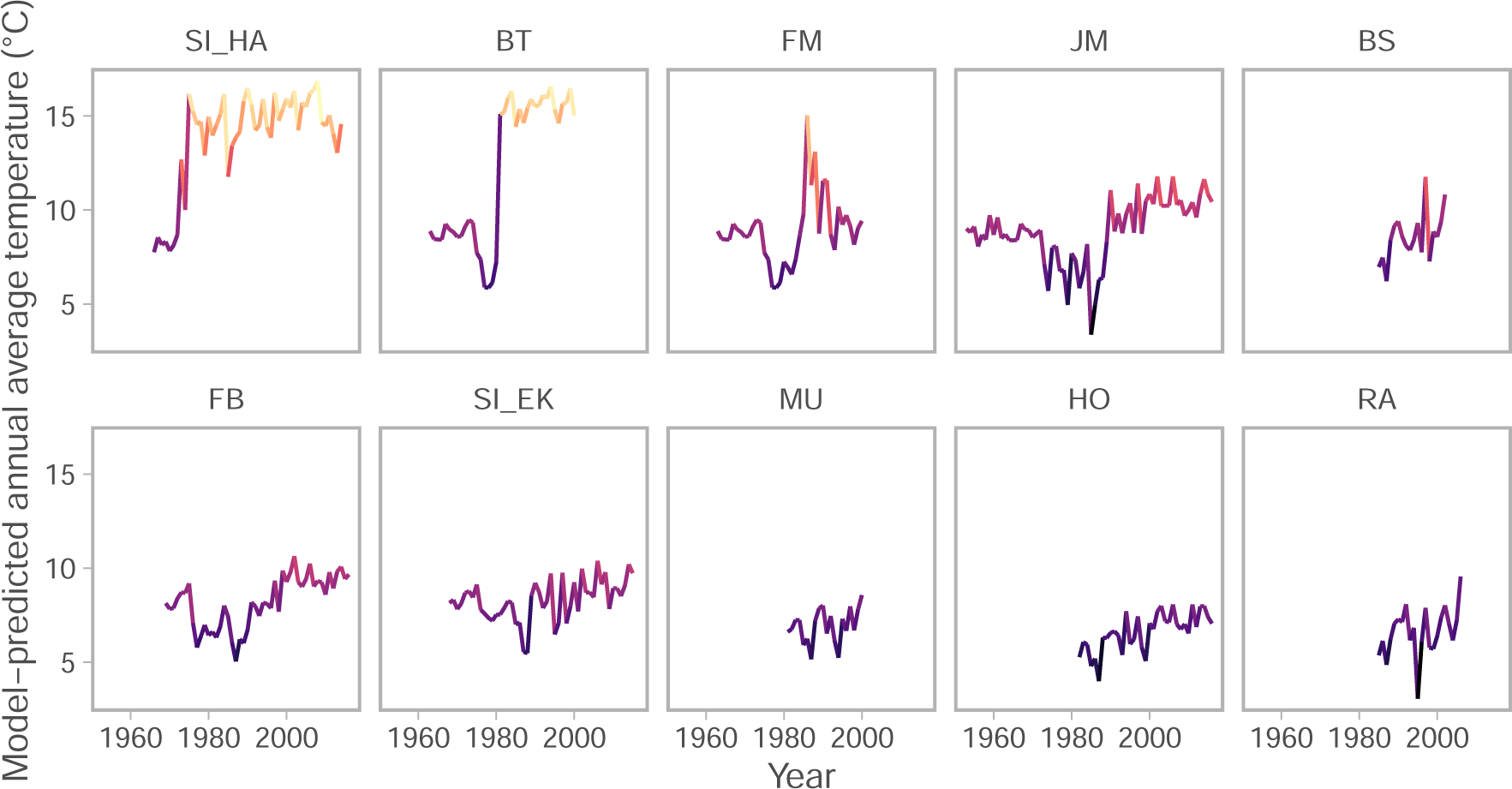
Annual average sea surface temperature as predicted by the GAM-model fitted to three temperature sources. Colour indicates temperature. Areas SI HA and BT have been heated by warm water discharge from nuclear power plants since 1972 and 1980, respectively.

We reconstructed local temperatures at each fishing site using three types of temperature data: automatic temperature loggers deployed near the fishing sites, manually measured temperatures at the time of fishing, and extended reconstructed sea surface temperature, ERRST (Huang *et al*. 2017). We chose these three types because they are complementary. Logger data provide daily temperatures during the ice-free season but do not go back as far in time as the growth data. Temperatures at the fishing event give a snapshot temperature at the site, and go back as far in time as we have fishing data. However, temperatures during fishing may not be representative of the whole growth season, and since we work with back-calculated length-at-age, we also need temperatures for years prior to fishing. Therefore, we also used modelled temperature time series (ERRST), which both provide good seasonal coverage and extend far back in time, but have a much coarser spatial resolution than the other sources. These three temperature data sources overlap in time (Supporting Information S2, Fig. S5), which allowed us to standardize the data using a statistical model (see next section).

### Statistical analyses

Individual-level growth models

To characterise individual growth rates, we fit von Bertalanffy growth equations — a special case of a Pütter model (Pütter 1920). The model describes growth rate in weight *w* as the difference between the rates of energy input (or anabolism) and energy expenditures (or catabolism):

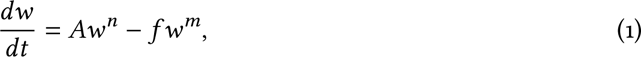

where *A* and *f* are the coefficients for anabolism and catabolism, respectively, and *n* is the size scaling of anabolism (von Bertalanffy 1938). With the assumption that catabolism is proportional to *w* (*m* = 1) and that *w* = *aL*^*b*^, where *a* is a condition factor and *L* is length, it can be integrated to the following form (Essington *et al*. 2001):

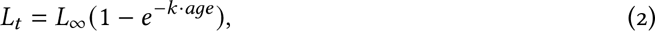

where *L*_*t*_ is the size (mm) at age *t* (years), *L*_∞_ the asymptotic size (mm), and *k* is the growth rate coefficient (year^−1^). It is however not a growth rate *per se* (which has unit size per time), but is instead related to the time it takes to reach the asymptotic size. Following van Denderen *et al*. (2020) and Andersen (2019), we further assume that the condition factor *a* is constant such that we can acquire *A* from equation (1) as *A* = 0.65*kL*_∞_. *A*, in contrast to *k*, can be interpreted as a size-corrected growth coefficient. Henceforth we refer to *A* this as the growth coefficient, and *k* in equation 2 to simply *k*.

We fit equation 2 to the multiple observations of back-calculated length-at-age for each individual using non-linear least squares (nls function in R (R Core Team 2020), version 4.2.3). We only used length-at-age, meaning only length at a back-calculated integer age (i.e., length at the formation of the age-ring), because sampling has occurred in different times of the year. We fit this model to every individual age 5 or older to ensure enough data points per individual to reliably fit the model. The filtering resulted in 142 023 data points across 23 605 individuals. We then calculated the median *k* by cohort and site across individuals (resulting in *n* = 306 *k* values) (Fig. 2).

### Model-based standardisation of local temperatures

In order to relate the site- and cohort-specific growth coefficients to temperature over time, we reconstructed average annual temperature sea surface temperature (*sst*) for each site using generalized additive models assuming Student-t distributed residuals to account for extreme observations:

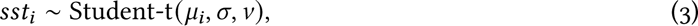

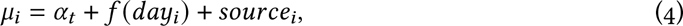

where *μ*_*i*_ is the mean *sst*, *σ* is the scale and *v* is the degrees of freedom parameter. *v* was not estimated within the model, but found by iteratively testing different values and visually inspecting QQ-plots to see how well the model could capture the heavy tails in the data. We used two sets of values, *v* = 6 for sites BS (Brunskär), BT (Biotest), FB (Finbo), FM (Forsmark), MU (Muskö), RA (Råneå) and SI EK (Simpevark Ekö) and *v* = 10 for HO (Holmön), JM (Kvädöfjärden), and SI HA (Simpevarp Hamnefjärden) (Supporting Information S2, Fig. S6). The parameter *α*_*t*_ is the mean *sst* of year *t* (included as factor), *f* (*day*) is a global smooth implemented as a penalized cyclic spline (i.e., the ends match—in this case December 31^st^ and January 1^st^) for the effect of day-of-the-year, and source is the mean temperature for each temperature source. We fit the temperature models by site separately, because the presence of artificial heating from nuclear power plants warranted complicated interactions between time, source and site in a global model, and those models did not converge. We fit our models in R using the package sdmTMB (Anderson *et al*. 2024, 2021) (version 0.3.0.9002), which uses mgcv (Wood 2017) to implement penalized smooths as random effects, and TMB (Kristensen *et al*. 2016) to estimate parameters via maximum marginal likelihood and the Laplace approximation to integrate over random effects.

We assessed convergence by confirming that the maximum absolute gradient with respect to all fixed effects was *<* 0.001 and that the Hessian matrix was positive-definite. We evaluated consistency of the model with the data by visually inspecting QQ-plots of randomized quantile residuals (Dunn and Smyth 1996) with fixed effects held at their maximum likelihood estimates and random effects sampled via Markov chain Monte Carlo (MCMC) (Thygesen *et al*. 2017) using Stan (Carpenter *et al*. 2017, Stan Development Team 2024) via tmbstan (Monnahan and Kristensen 2018)

### Effects of temperature on growth coefficients

To estimate how von Bertalanffy growth parameters were related to temperature we fit Bayesian generalized mixed models with site-varying and correlated intercepts and slopes and Student-t distributed residuals to account for extreme observations:

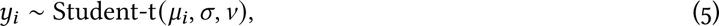

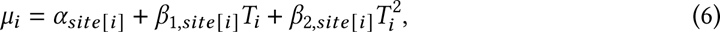

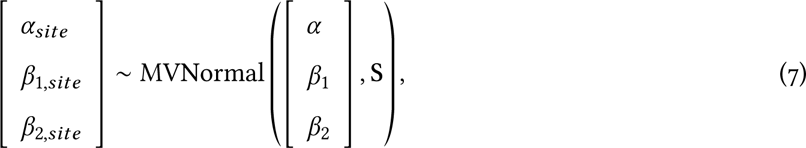

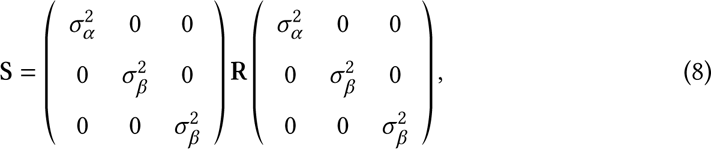

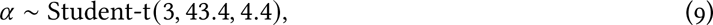

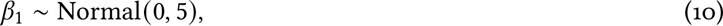

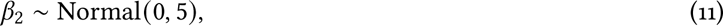

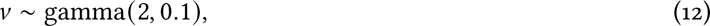

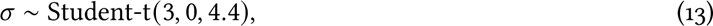

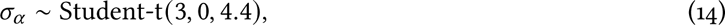

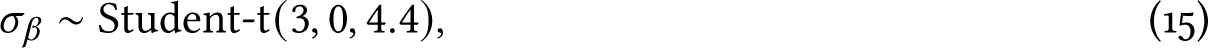

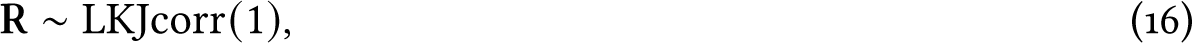

where *μ* is the mean, *σ* is the scale and *v* is the degrees of freedom parameter. We scaled temperature, *T*, by subtracting the mean and dividing by the standard deviation before squaring to reduce the correlation between the two variables (Schielzeth 2010). *α* is the intercept and *β*_1_ and *β*_2_ are the coefficients for temperature and temperature squared. S the covariance matrix with the correlation matrix, R, of site random effects and their correlations factored out. We used the default prior specification for the overall intercept as implemented in brms, and a Normal(0, 5) for regression coefficients. The *σ* priors had a lower bound of 0. The prior for the correlation matrix R is a LKJcorr(1), which is flat and puts similar probability on all possible correlation values (correlation between random effects), *ρ*. We fit alternative models with other fixed and random effects (e.g., only random intercepts and linear temperature effects only). A full overview of the alternative models and their expected predictive accuracy (expected log pointwise predictive density, elpd) using Pareto smoothed importance sampling to approximate leave-one-out cross-validation (Vehtari *et al*. 2017) can be found in Supporting Information S1, Table S1. The model presented here (equations 5–16) was selected because it has a good fit to data, and its elpd was not substantially different from the model with the highest elpd, and is complex enough to allow site-specific and non-linear temperature responses (Gelman *et al*. 2021). We fit the same model to von Bertalanffy growth coefficients *k* and *L*_∞_, with the modification that the prior for the intercept *α* was Student-t(3, 0.2, 2.5) and Student-t(3, 340.9, 80), respectively.

We fit the model using Stan (Carpenter *et al*. 2017, Stan Development Team 2024) via the R package brms (Bürkner 2017, 2018) (version 2.20.4). We sampled from the models with 4000 iterations each on 4 chains, discarding the first 2000 as warmup. Model convergence and fit were assessed by ensuring potential scale reduction factors were factors were smaller than 1.01 which suggests all four chains converged to a common distribution, Gelman *et al*. (2003), as well as by visually inspecting posterior predictive checks (Supporting Information S1, Figs. S2, and S4). Bayesian *R*^2^ values were calculated using the R package performance (Lüdecke *et al*. 2021), which implements the method described in Gelman *et al*. (2019). We made conditional predictions and manipulated posterior draws with the R package tidybayes (Kay 2019).

## Results

We find large inter-annual fluctuations in annual average temperatures between sites, and increasing trends over time in some sites (Fig. 3). Due to the spatial and temporal range of data, and the artificial heating from nuclear power plant water discharges, we observe large contrasts in average temperatures, which were not clearly related to latitude (Fig. 2). Across all sites, mean annual average temperatures range from 3°C–17°C, and the largest range within a site (over time) is 6°C– 16°C (site BT). Individual growth trajectories of fish showed large variation within and across sites (Fig. 4). Site-specific growth coefficients (*A*) generally increased over time, but not always linearly and not synchronously across all sites, indicating that local drivers shape variation in growth between cohorts (Fig. 2).

**Figure 4:**
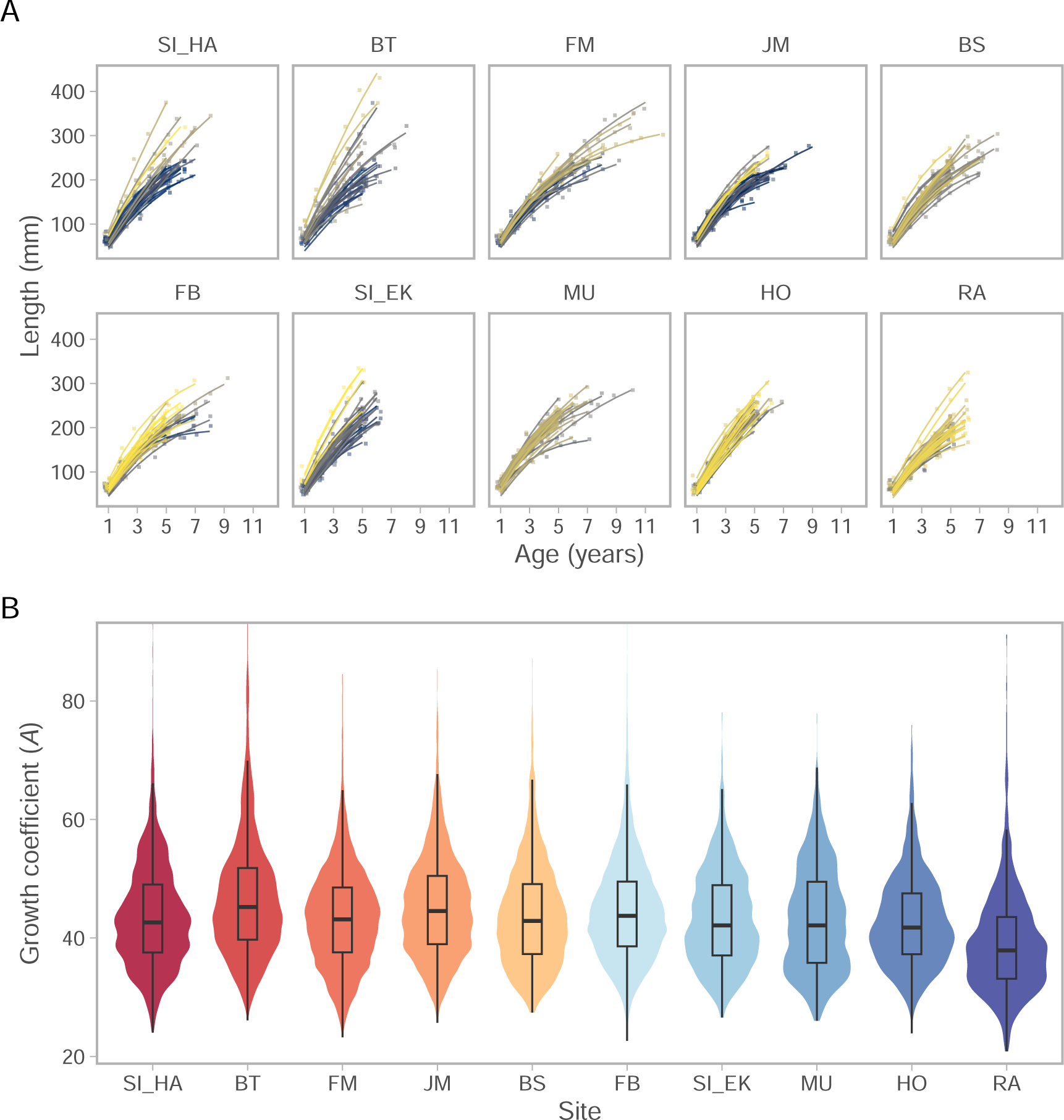
Length plotted against age for all sites (A). Points are data for 30 randomly selected individuals (indicated by colour) in each site, and lines are the predicted von Bertalanffy growth curve. Panel B depicts the distribution of von Bertalanffy growth coefficients *A*, where colours are based on the mean temperature across all years, as violins, and quantiles depicted as boxplots.

Models relating the growth coefficient *A* to temperature that allowed for site-specific estimates (as temperature-site interactions or site-varying parameters) were indistinguishable based on the leave-one-out cross-validation, as the expected log predictive density (Δ*elpd*) was less than four (Supporting Information S1, Table S1) (Sivula *et al*. 2023). We therefore used the model with site-varying intercepts and site-varying effects of temperature and allowed for non-linear temperature relationships by using linear and quadratic temperature terms. The variance explained by the model (Bayesian *R*^2^) with fixed and random effects was 0.26. The distribution of cohort and site-specific growth coefficients *A*, was characterized by heavy tails compared to a Gaussian distribution, and the *v* parameter (equation 5) was estimated to 5.9. Predictions from the full model revealed that growth coefficients increased with temperature initially in all sites. However, the growth coefficients in two of the three warmest sites either plateaued or declined with warming at high temperatures (sites FM and SI HA in Fig. 5).

**Figure 5:**
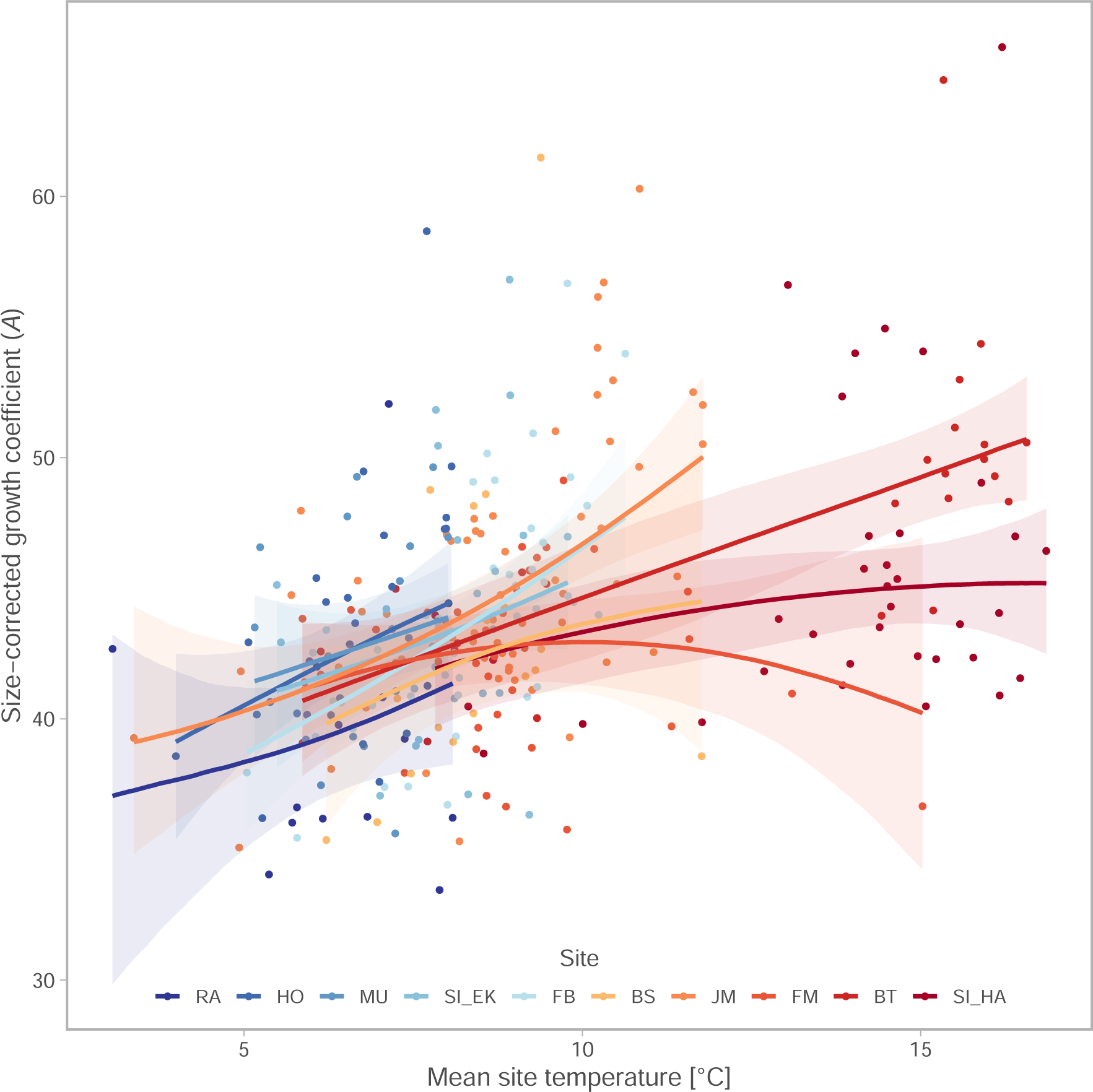
von Bertalanffy growth coefficients, *A*, as a function of temperature. Each point depicts the median growth coefficient for a cohort and site, and the coloured lines depict the median of draws from the expectation of the posterior predictive distribution and ribbons the 90% credible interval from the linear mixed effect model for each site.

Population-specific growth-temperature curves mapped closely onto a pooled ‘global’ growth-temperature curve across all populations, and site-specific growth coefficients (random intercepts) were not related to the average site temperature (Fig. 6). This pattern was also found for the von Bertalanffy growth parameters *k* and *L*_∞_ (Supporting Information S1, Fig. S1.

**Figure 6:**
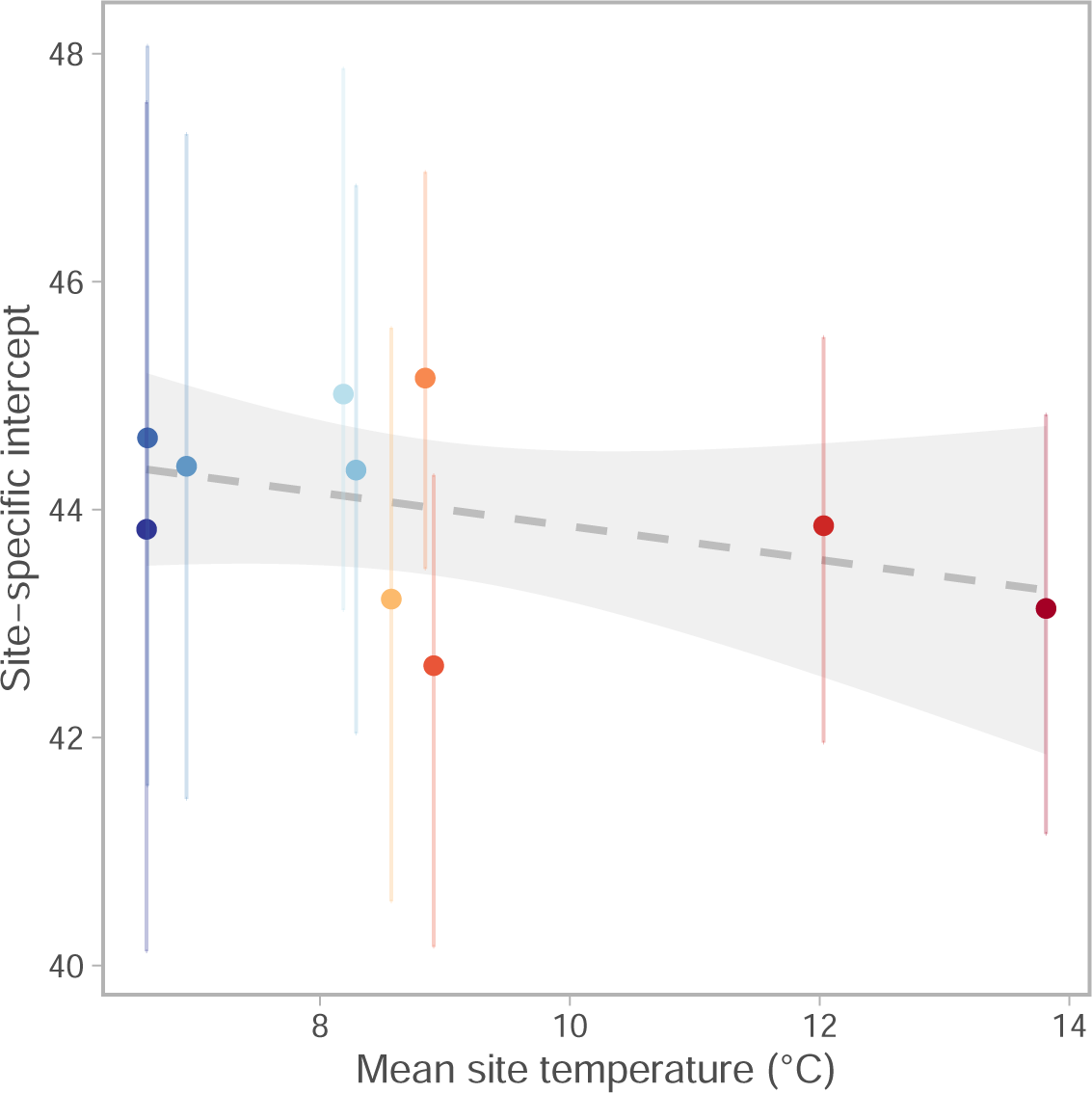
Site-specific intercepts, which corresponds to the growth coefficient at average cohort temperature across all sites, as a function of the mean site temperature. The line and ribbon depict the fit of a linear regression (slope is not significant).

## Discussion

Our findings indicate a lack of local adaption in growth to local temperatures, despite differences in experienced environmental temperatures and reproductive isolation among populations. Populations are expected to grow similarly at the average temperature irrespective of origin (Fig. 6), i.e., populations grow similarly where their experienced temperature ranges overlap (Fig. 5). If populations had adapted to local temperatures, we expect similar effect of warming across populations, assuming they occupy temperatures slightly below the local optimum temperature (Ohlberger 2013). Instead, we find that populations in relatively cold environments will benefit from climate warming via increased body growth rates up to a certain ‘global’ temperature optimum, whereas populations in relatively warm environments will experience reduced growth due to the negative effects of warming beyond their optimum growth temperature.

In line with our results, Neuheimer *et al*. (2011) found that for populations of banded morwong (*Cheilodactylus spectabilis*), increasing temperatures were associated with reduced growth rates for the population at the warm edge of the species’ distribution (New Zealand) but higher growth rates for populations at the colder edge of the range (Tasmania). Similarly, Morrongiello and Thresher (2015) found that body growth of tiger flathead in populations off Southeast Australia increased with temperature but not in the warmest area. In terms of body size, Beaudry-Sylvestre *et al*. (2024) recently found that the size of four-year old Atlantic herring (*Clupea harengus*) in the Northwest Atlantic followed a similar pattern — populations in warmer regions tended to have negative associations with of warming. Analogously, English *et al*. (2022) found that groundfish in the Northeast Pacific often responded positively to warming if they were in cool locations, and negatively if they were in warm locations (where both biomass and temperature change were expressed as velocities). Collectively, these and our findings are in contrast to the common finding in invertebrate species that exhibit local adaptation in growth to temperature, even over small spatial scales (Sanford and Kelly 2011). These results illustrate the importance of testing for population-specific temperature-sensitivities when studying species responses to warming, and of accounting for both the rate of climate change and baseline temperature conditions.

The ability to adapt to local environmental conditions allows populations to expand their range and better cope with spatially varying environmental conditions (Kirkpatrick and Barton 1997). Changes in trait-temperature relationships due to thermal adaptation in natural populations are expected in response to climate warming (Angilletta 2009), and previous studies have shown that local adaptation in physiological traits can facilitate different thermal optima among populations (e.g., Atlantic cod (Righton *et al*. 2010), kelp (Britton *et al*. 2024), corals (Howells *et al*. 2013) and invertebrates (Sanford and Kelly 2011)). However, adaptive capacities and the pace of thermal adaptation differ among species (Martin *et al*. 2023) and depend on life-history trade-offs, underlying genetic variation, the potential for gene flow (Kirkpatrick and Barton 1997), and environmental conditions. The apparent lack of contemporary thermal adaptation in Baltic Sea perch, despite low gene flow between populations due to limited dispersal and movement (Bergek and Björklund 2009), indicates limitations in evolutionary changes to local temperature. This suggests that similar factors may also limit future thermal adaptation that would allow local populations to better withstand changing temperatures. A low adaptive capacity implies that body growth rates in populations already experiencing temperatures around or above their thermal optimum will decline with further warming. This will likely result in lower biomass production in warm environments, as observed, for example, across spatial temperature gradients (van Dorst *et al*. 2019).

Our study also illustrates the importance of accounting for unimodal temperature dependencies. Often simpler models like the exponential Arrhenius equation are used to model biological and ecological processes (e.g., Savage *et al*. (2004), Vasseur and McCann (2005), Lindmark *et al*. (2018)), under the assumption that the ‘biologically relevant temperature range’ which species occupy is below their optimum. Growth rates are only exponentially related to temperature even further from the optimum, i.e., below the inflection point of the unimodal curve. We find this supra linear temperature curve in only two of the cooler sites. Over the entire temperature range, the temperature curves tend to flatten out even though a true optimum curve is only found in one population (Fig. 5), hence, temperatures close to or above the optimum are therefore biologically relevant, in which case models other than the Arrhenius equation are more appropriate.

There are a number of limitations to our analysis. For instance, growth in temperate regions varies over the year and no single temperature metric may fully reflect thermal conditions that determine cohort-specific growth rates. Given that growing season lengths differ in our data set due to different light conditions, we opted to use a simple annual average. Degree days (the integral of time above a certain temperature threshold) is an often recommended metric (Neuheimer and Grønkjær 2012), but there is some uncertainty in temperatures below which growth does not occur, even for a well studied species like perch (Karås and Thoresson 1992), how starvation during cold periods affects growth trajectories, and whether or not that varies between sites. Lastly, it is not straightforward to formally test for differences in thermal optima between populations because populations generally occupy temperatures below their optimum.

## Conclusion

Our findings suggest that mean environmental temperatures during warm years have reached or surpassed the optimum growth temperature for two of the examined populations (Fig. 5), but that most populations have a positive, linear relationship with temperature. Our ability to detect this pattern relies heavily on the length of the time series as well as the unusually large temperature contrasts due to warm water pollution from nuclear power plants, which highlights the importance of long term environmental monitoring across environmental gradients. Considering the lack of evidence for local adaptation to temperature, we expect that adverse effects of continued warming on Baltic Sea perch will accumulate and decrease individual growth rates in the warmest populations. Similar constraints on adaptive capacities in response to warming can be expected for other species of fish, and ectotherms more generally (Pawar *et al*. 2024).

## Acknowledgements

We are very grateful to all the staff involved in data collection over the years and to the funding bodies of the long-term monitoring. The study was financed by the Swedish Research Council Formas (grant no. 2022-01511 to Max Lindmark) and SLU Quantitative Fish- and Fisheries Ecology.

## Author contributions

All authors contributed to the manuscript. A.G. conceived the study, M.L. and A.G. prepared the raw data and J.O. contributed to preparing temperature data. J.O. and M.L. led the design with contributions from AG. M.L and J.O conducted the statistical analyses. M.L. wrote the first draft. All authors conceptualized the results, contributed to revisions, and gave final approval for publication.

## Data Availability Statement

All code and data to reproduce the results are available on GitHub (https://github.com/maxlindmark/ perch-growth), and will be deposited on Zenodo before publication.

