## supporting information for "Stronger effect of temperature on body growth in cool than in warm populations suggests lack of local adaptation"

### S1. von Bertalanffy model

Table S1: Δelpd (expected log pointwise predictive density relative to the best fitting model) for all models relating $A$ to temperature. The difference between models in bold is considered negligible (Sivula et al., 2023)

| Model description | Δelpd | s.e. of Δelpd |
|---|---|---|
| **Interaction between temperature and site, no random effects** | 0 | 0 |
| **Random intercept and slope, linear temperature effects** | -0.2 | 5.1 |
| **Interaction between area and temperature, common squared term, no random effects** | -0.8 | 0.5 |
| **Linear and quadratic temperature effects, full random effects** | -0.9 | 2.4 |
| Linear and quadratic temperature effects, no interactions nor random effects | -6.1 | 4.1 |

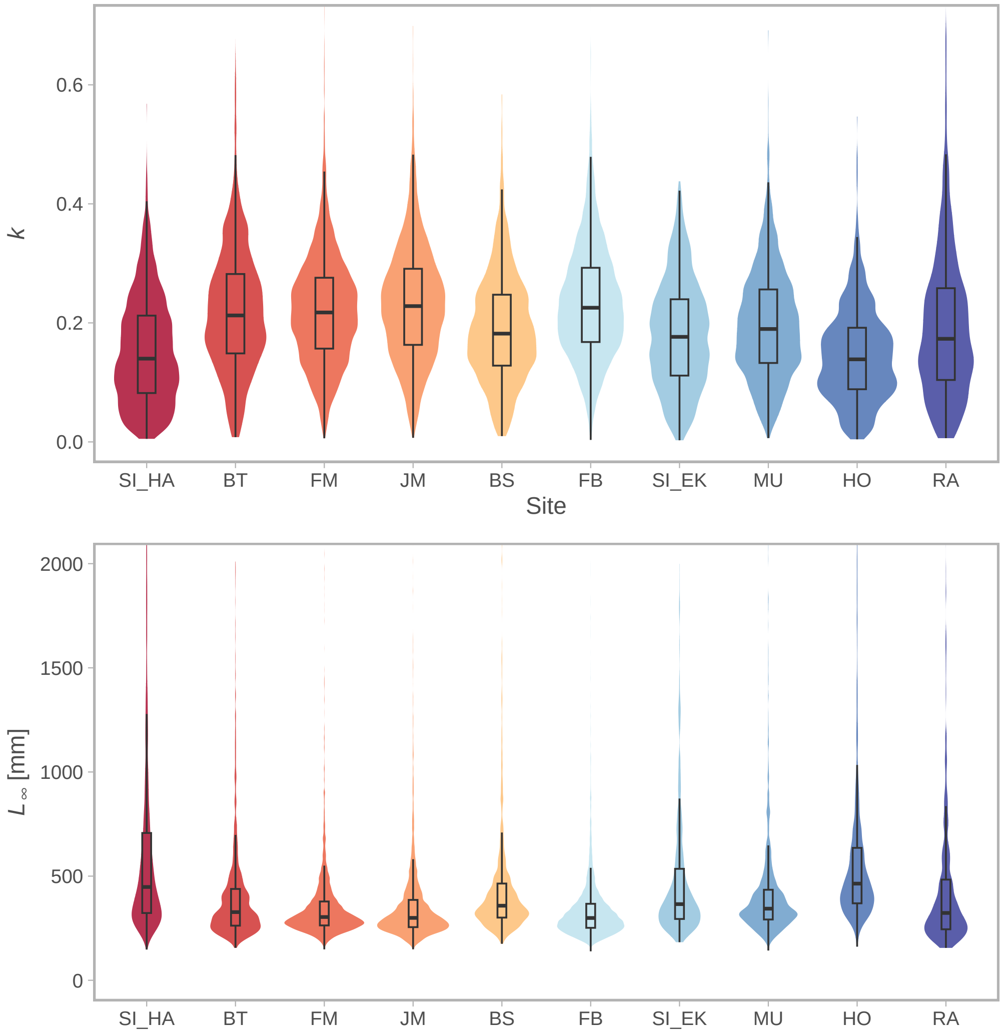


Figure S1: Distribution of von Bertalanffy growth parameters $k$ (top) and $L_\infty$ (bottom), where colours are based on the mean temperature across all years, as violins, and quantiles depicted as boxplots.

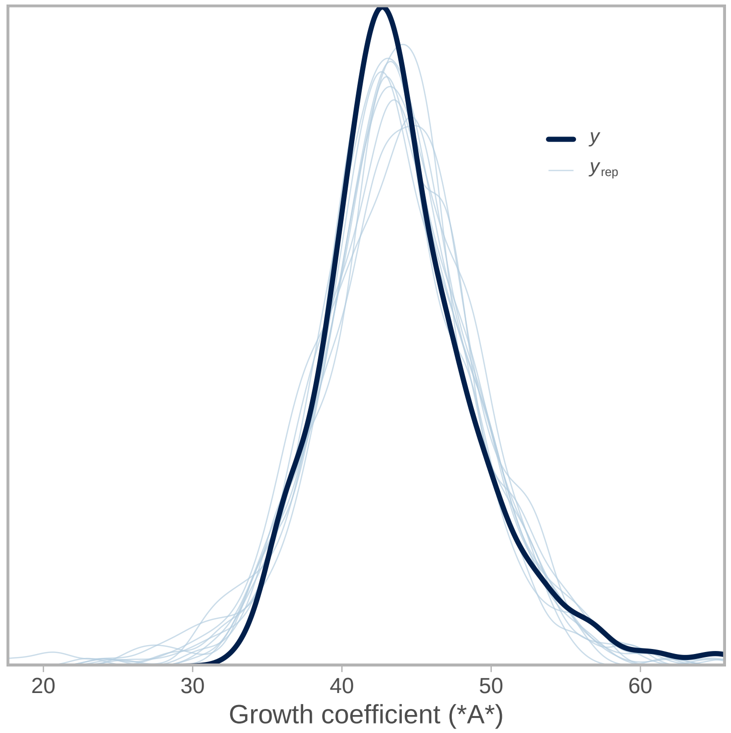


Figure S2: Graphical posterior predictive checks depicting the distribution of data (dark blue) in relation to 10 sampled datasets (light blue).

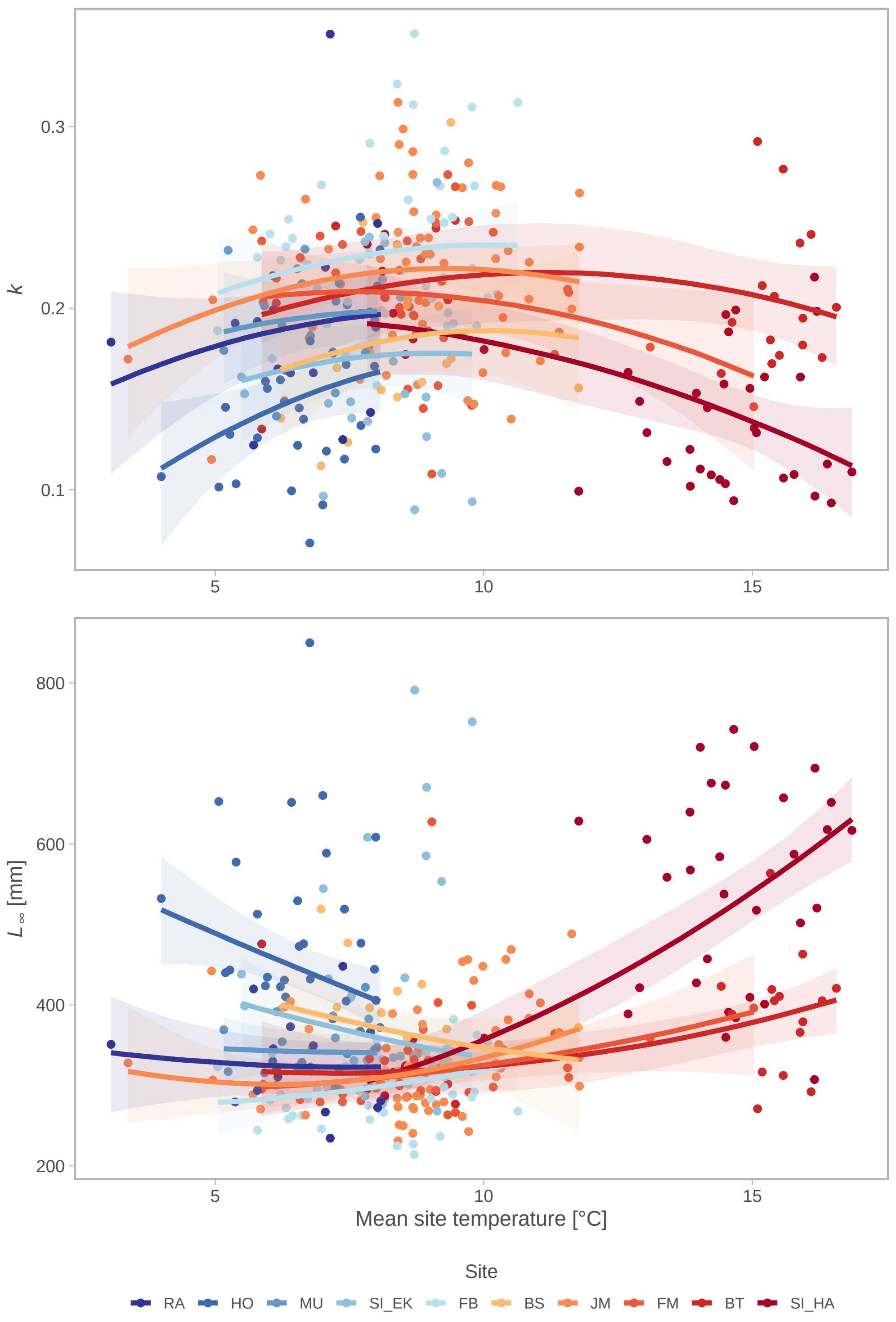


Figure S3: von Bertalanffy growth coefficients $k$ (top) and $L_\infty$ (bottom), as a function of temperature. In panel A, each point depicts the median growth coefficient for a cohort and site, and the coloured lines depict the median of draws from the expectation of the posterior predictive distribution and ribbons the 90% credible interval from the linear mixed effect model for each site.

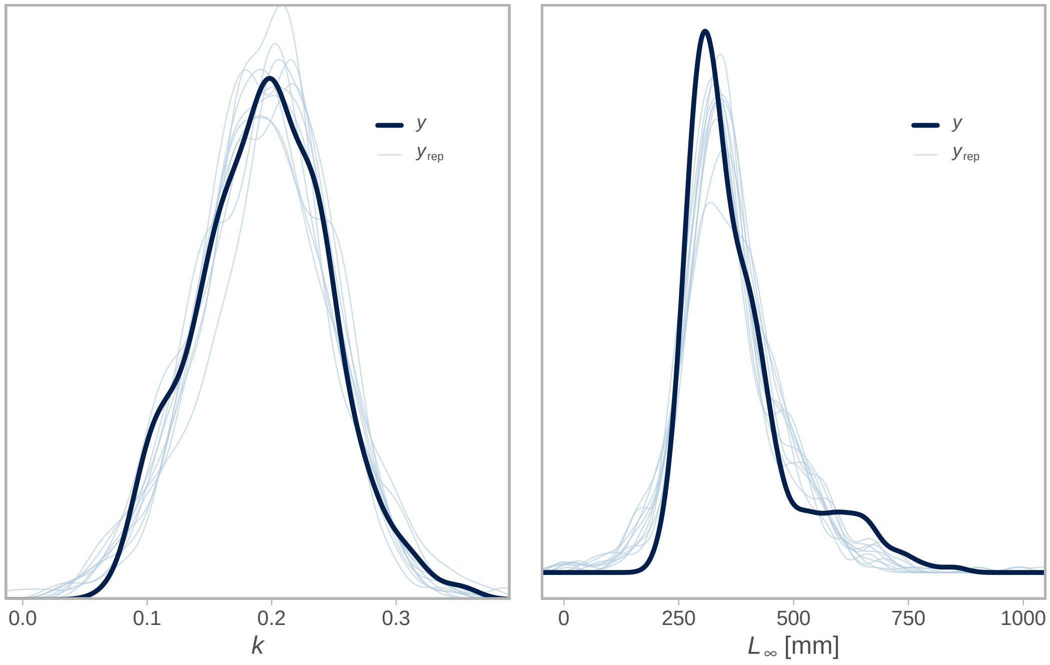


Figure S4: Graphical posterior predictive checks depicting the distribution of data (dark blue) in relation to 10 sampled datasets (light blue) for $k$ (left) and $L_\infty$ (right).

### S2. Temperature model

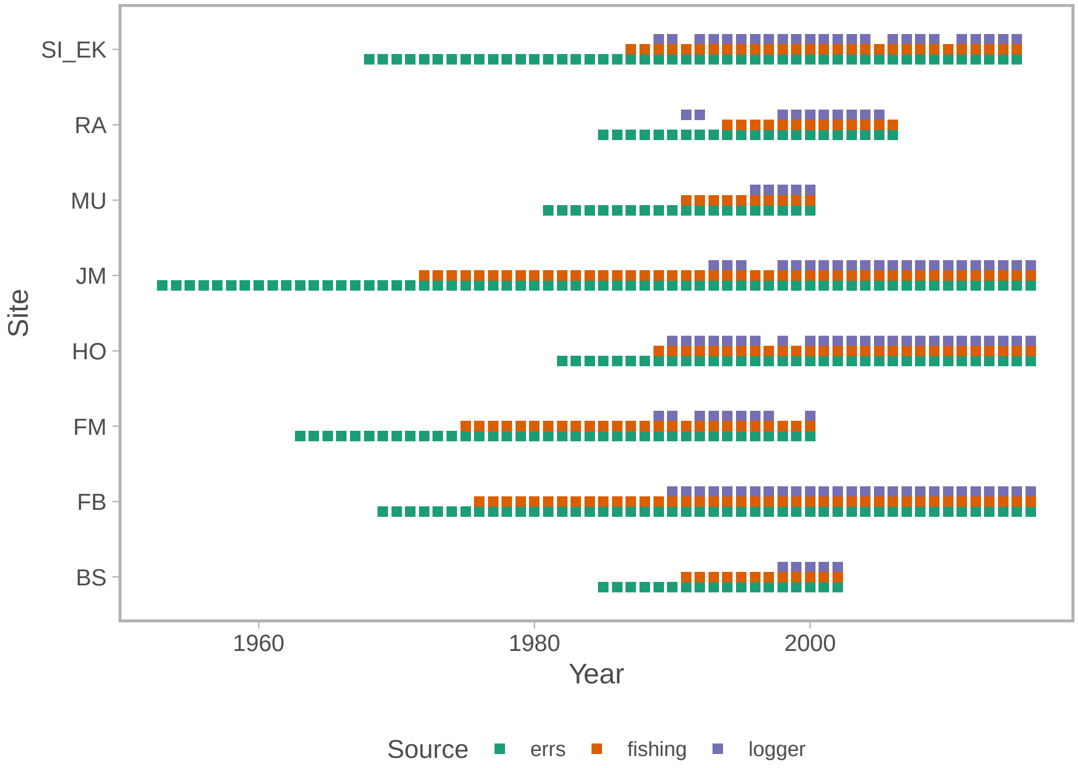


Figure S5: Available temperature data by year, site and source.

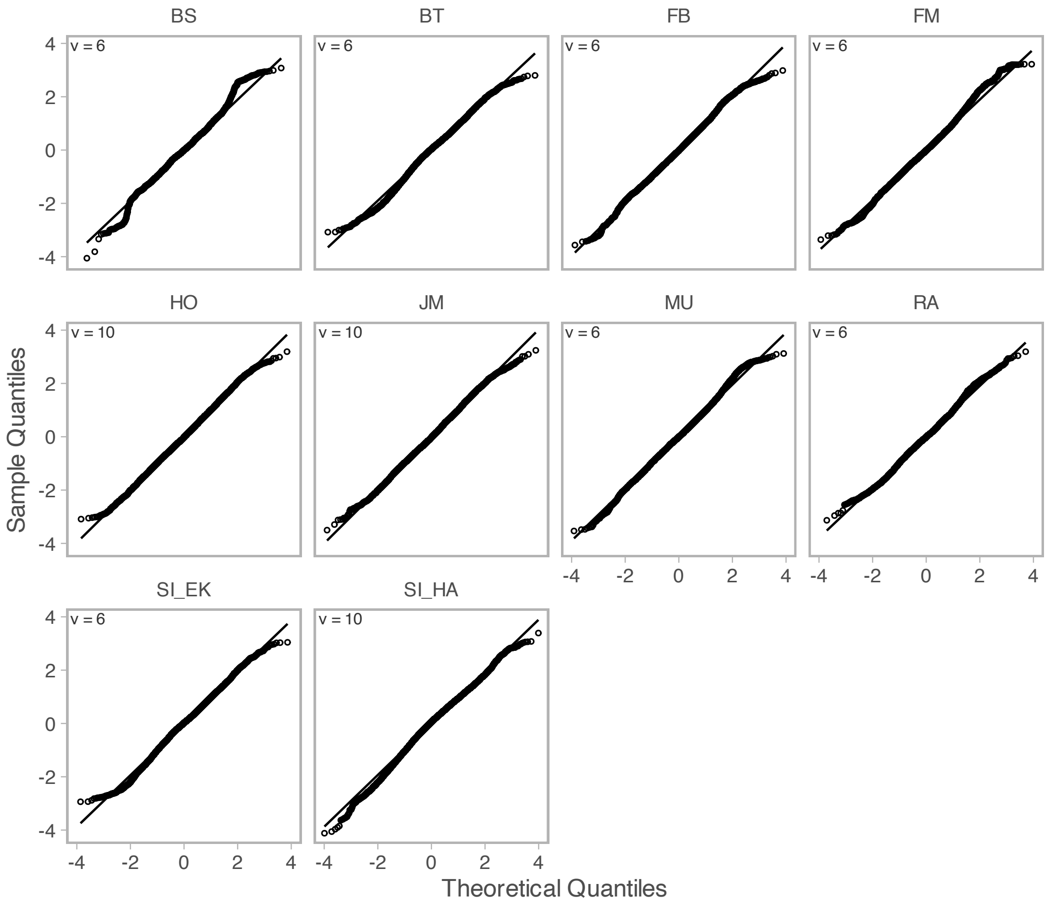


Figure S6: QQ-plot of site-specific temperature models based on randomized quantile residuals, where fixed effects are held at their maximum likelihood estimate and the random effects are sampled with MCMC via `tmbstan` (Monnahan and Kristensen, 2018) and `Stan` (Stan Development Team, 2022), in line with (Thygesen et al., 2017; Rufener et al., 2021). The degrees of freedom parameter is printed in the top-left corner.

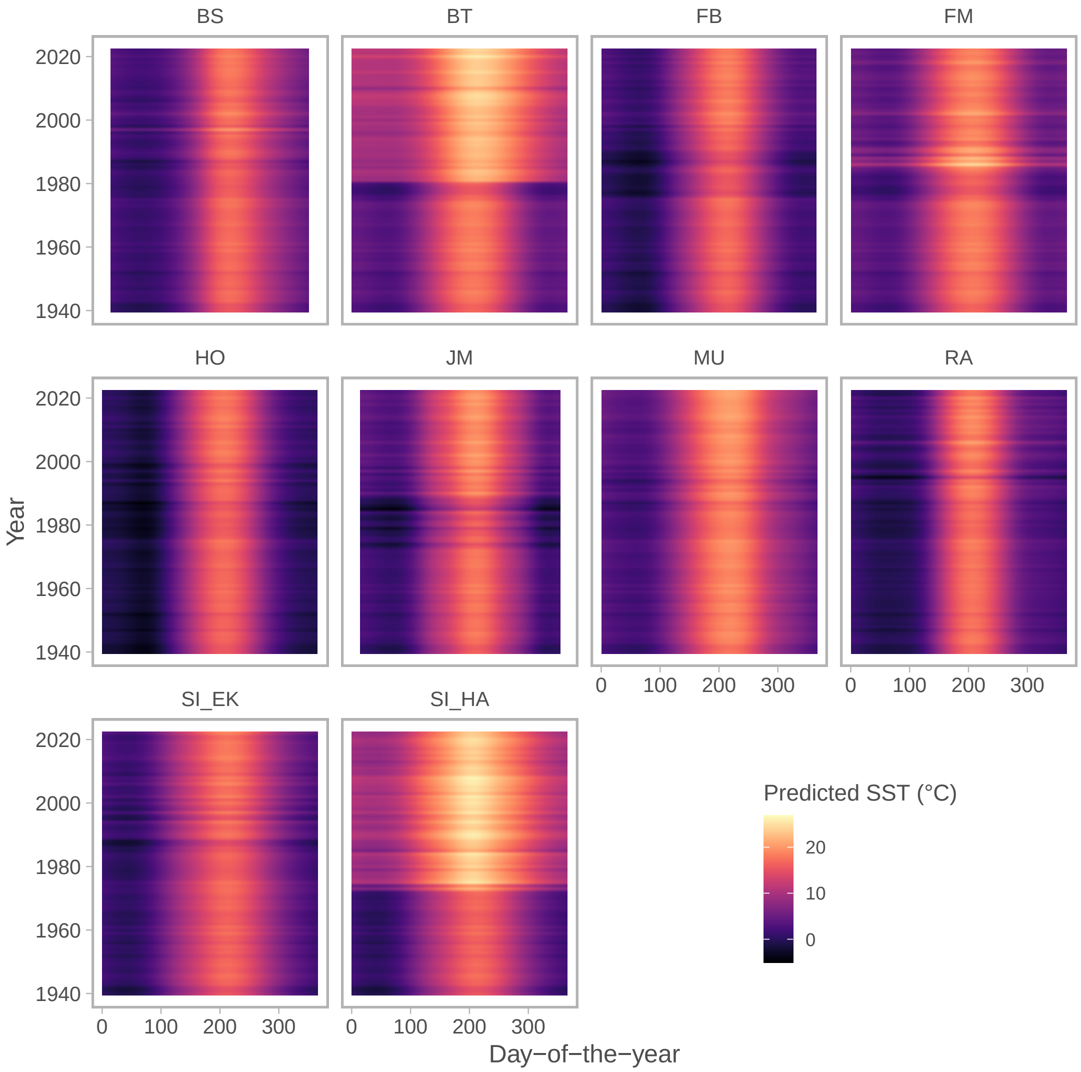


Figure S7: Predicted temperatures by site, year (y-axis) and day of the year (x-axis), where the color indicates the temperature. Predicted at the logger level.

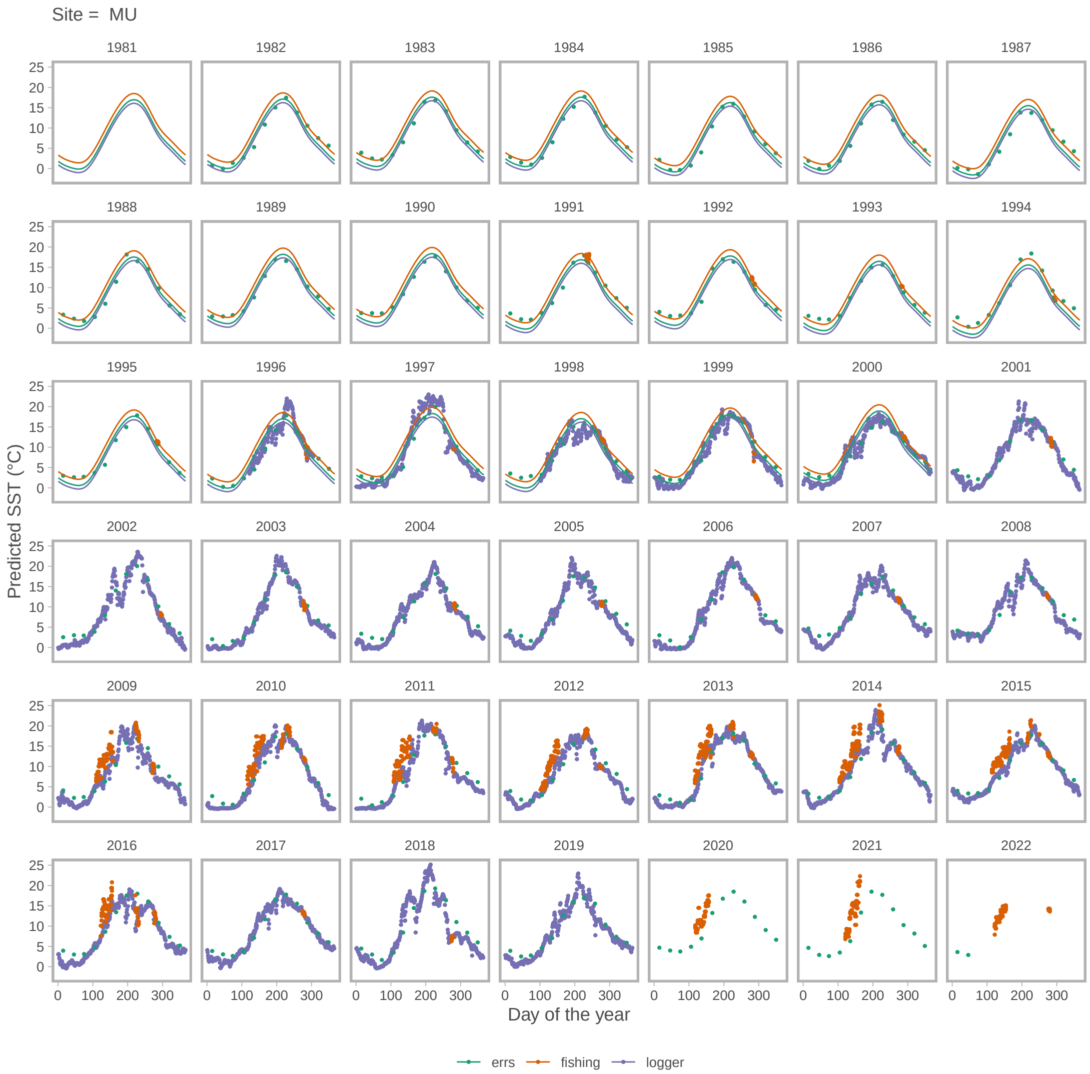


Figure S8: Predicted temperatures (y-axis) by day of the year (x-axis) for all three temperature sources, indicated by color, using Muskö (MU) as an example.
